# Pervasive effect of bottom trawling on benthic biodiversity unravelled by metabarcoding across the Tree of Life

**DOI:** 10.64898/2026.10.05.756630

**Authors:** Chloé Beaulieu, Sandrine Vaz, Thomas Hénault, Céline Labrune, Christine Barras, Magali Schweizer, Jadwiga Rzeznik-Orignac, Grégoire Maillet, Audrey Pruski, Nicolas Henry, Sophie Arnaud-Haond

## Abstract

Bottom trawling is the most widespread human disturbance on the seafloor, known to affect seabed structure and benthic biota. Yet, its impact on benthic communities remains documented mainly for taxa identifiable through morphology, leaving microscopic benthic communities largely overlooked. The Gulf of Lion continental shelf has been trawled for decades by French and Spanish fleets, to the point that encountering untrawled reference areas became elusive.

Here, we examined prokaryote, protist and Animalia benthic communities at eight stations, using eDNA metabarcoding of core sediment samples, with five primer-pairs. Benthic communities were cross-referenced with contrasted trawling impacts, integrated across three periods: over one month, three years or ten years preceding the sampling.

The community composition was primarily structured by ecological habitats, yet, trawling pressure showed association with prokaryotes and Animalia diversity. Diversity was higher at sites trawled more intensively in the preceding month, whereas long-term trawling was associated with a significant erosion of Animalia alpha diversity. Unexpectedly, communities did not display consistent vertical structure and the relative contributions of trawling and hydrodynamic reworking remain unresolved due to the absence of untrawled sites.

Overall, our results show that trawling effects extend to prokaryotic compartments rarely considered in impact assessments, providing a proof of concept for eDNA-based assessment, and underline a pervasive long-term effect on Animalia communities. This study also highlights the importance of the timescale used to quantify fishing pressure and underscores the complexity of impact assessment when monitoring begins too late to establish a clear baseline, particularly when no reference site remains.

## 1. Introduction

Among human activities that impact the seafloor, bottom trawling is the most widespread geographically (Halpern et al., 2008; Hiddink et al., 2017). The rise of industrial fishing in the mid-20th century was accompanied by improvements in the efficiency of many methods, particularly bottom trawling methods, whose footprint has gradually increased, in conjunction with the depletion of coastal stocks (Amoroso et al., 2018).

Bottom-trawling impact on the ocean floor has been demonstrated physically through sediment resuspension and habitat homogenization (Pilskaln et al., 1998; Puig et al., 2012), geochemically, notably through alterations in carbon, nitrogen and silicate budgets (Olsgard et al., 2008; Pusceddu et al., 2005); and biologically, through changes in the size distribution of living organisms, the abrasion of biogenic structures (Schwinghamer et al., 1998), the loss of keystone species (Buhl-Mortensen et al., 2016), and a reduction in total biomass or species richness (Hiddink et al., 2017, 2006). The latter is all the more significant when trawling occurs in areas subject to low natural disturbance and in deep waters (Hiddink et al., 2006; Queirós et al., 2006; Pusceddu et al., 2014; Leduc et al., 2025). The combined effect of these alterations reaches such levels that it has been compared to forest clear-cutting in the terrestrial environment (Watling and Norse, 1998).

The first large-scale demonstration of the effect of trawling on benthic fauna showed that trawled areas supported much smaller organisms, with a decline in faunal production due to the loss of larger organisms (Jennings et al., 2001). These results obtained in the North Sea are similar to those later reported in the Bay of Biscay at depths of about 100 m, where areas under moderate fishing pressure already seem to support only a handful of fish and invertebrate species, but those under heavy fishing pressure appeared completely depleted (Blanchard et al., 2004). Furthermore, the key role of bioturbators for benthic ecosystems and macrofauna has been demonstrated experimentally (Widdicombe et al., 2004), and their sensitivity to trawling therefore suggests the existence of a cascade effect on the entire community.

The surface of highly fished areas may be swept entirely over 10 times per year (Eigaard et al., 2017; Hiddink et al., 2026); recovery is therefore often illusory, especially since in some deep-water areas signs of improvement are extremely weak, even after 15–19 years without exploitation (Clark et al., 2016; Goode et al., 2025). Conversely, areas protected from trawling, such as the Menorca Channel, demonstrate the effectiveness of these measures through the presence of numerous vulnerable species, ranging from sponges to gorgonians (Grinyó et al., 2018).

While the impact of trawling on the resource has been the subject of numerous studies, the paucity of knowledge regarding smaller size fractions of the benthic diversity, notably microbial communities (Seidel et al., 2026), prevents a global understanding of the extent of disturbances across different timescales, and may hinder the identification of biological indicators. Benthic communities play a major role in biogeochemical cycles, nutrient recycling, and ecosystem functioning, and bacterial and meiofaunal compartments are key components capable of responding rapidly to environmental disturbances due to their short generation times and/or metabolic reactivity (Giere, 2009; Murray, 2006). Although the sensitivity of small-sized meiofauna (nematodes and copepods) has been demonstrated through a combination of experimental and field approaches (Schratzberger et al., 2009; Schratzberger and Jennings, 2002), there is limited research currently available on these compartments, largely due to the logistical difficulties or even the impossibility of studying them using traditional morphological approaches or conventional culturing methods.

More recently, studies based on environmental DNA (eDNA) have made it possible to expand impact assessments to include those compartments invisible to classical optical enhancement techniques, and overcoming the observer effect to allow integrating studies conducted by specialists in different laboratories or geographic areas. They also allow for greater repeatability and standardization. Most studies conducted to date using this type of method have, however, focused on DNA present in water and vertebrate biodiversity (Jordan et al., 2026; Kasmi et al., 2024; Russo et al., 2021; Veron et al., 2023), even though some studies on benthic invertebrates (Good et al., 2022; Khodami et al., 2025; Oosthuizen et al., 2023) or microorganisms (Bradshaw et al., 2024; Seidel et al., 2026) start to emerge.

In the present study, we focused on eDNA contained in sediment in order to study benthic communities across the entire tree of life and compare their composition along an impact gradient. The aim is to assess the robustness of the method and its limitations, by i) testing for the hypothesis of a diachronic response to trawling of different organisms with contrasting life-history traits and ii) testing for the effect of different trawling pressures on physical habitats, notably on the layering of communities in the sediment. Sediment samples were collected in the western Mediterranean, in two areas of the Gulf of Lion, along a trawl impact gradient and analysed following an end to end (field to bioinformatic analysis) protocol developed by Arnaud-Haond et al. (2026). DNA from 5 layers of sediment depths down to 15 cm was subjected to metabarcoding of five representative fragments preferentially targeting prokaryotes (16S-V4V5), protists (18S-V9), and animals (COI & 18S-V1, V4 and V9). Ultimately, this work seeks to test for the ability of eDNA to help developing a more comprehensive and integrative framework for ecosystem-based fisheries management and conservation in trawled and protected seascapes.

## 2. Material and Methods

### 2.1. Area of study and sampling sites

The sampling took place in the continental shelf of the Gulf of Lion, in the north-Western part of the Mediterranean. This shelf is a gentle slope down to 90m, and drops off sharply into the deep sea due to the formation of submarine canyons, that can generate downwelling or upwelling. The Gulf of Lion is also influenced by the Liguro-Provençal current (that flows to the South-West), and submitted to river inflows (mostly from the Rhône River) that discharges sediments and organic matter into the oligotrophic Mediterranean Sea (Durrieu De Madron et al., 2000). Weather conditions (winds, heat) can also influence the water masses in a seasonal way, as does the summer thermocline, that creates a physical barrier in the water column around a 20 m depth (Millot, 1990). All these elements influence the dynamics of the water column, but also that of the benthic zone (Requena et al., 2013). Samples from 8 stations, located on the Gulf of Lion continental shelf, were collected in August 2021 during the IMPEC a cruise, conducted aboard the French R/V Europe (Labrune, 2021). Five stations are located near the head of the Lacaze-Duthiers canyon, around the 100 m isobath (area 1, stations A, C, E, G, J), while the 3 other stations were located in the middle of the continental shelf within the Gulf of Lion Marine Natural Park, on the 80 m isobath (area 2, stations K, O, P) (Figure 1).

**Figure 1:**
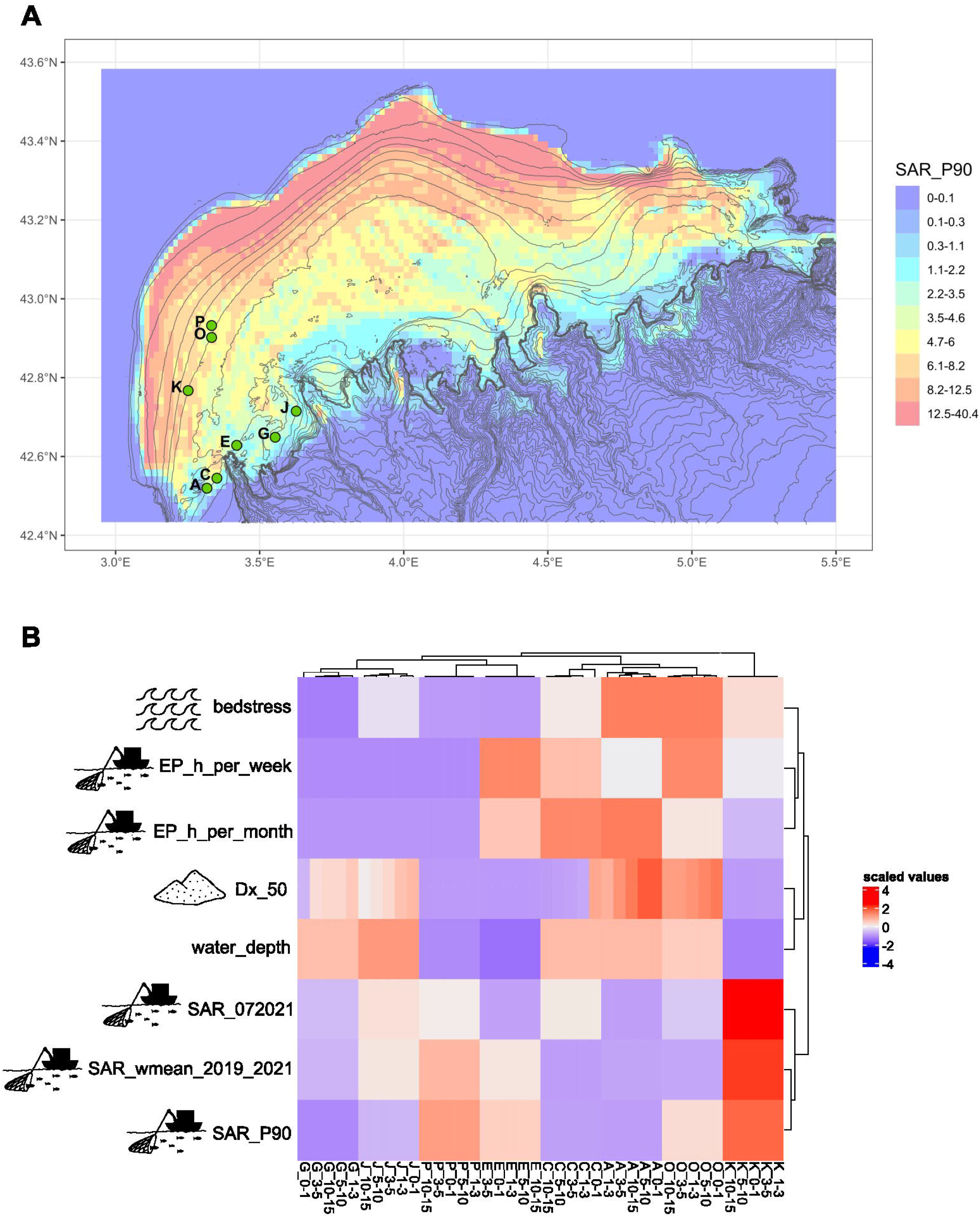
Information about the study sites. (A) Map of the Gulf of Lion, coloured depending on the abrasion of seabed due to trawling (90th percentile of SAR between 2012 and 2021: SAR_P90), with the localisation of study sites. (B) Heatmap of the values of the environmental and fishing variables at the different study sites. All the variables are scaled and centered for comparison (raw values available in the Supplementary Table 1). Red represents the highest values of the variable among the studied sites and blue the lowest values. Variables and stations are ordered by hierarchical clustering (euclidean distance with complete linkage clustering).

At each station, three replicate cores were collected from three separate deployment of the multicorer (Oktopus multicorer with 8 cores of 10 cm diameter and 30 cm length). The cores were sliced onboard in a lab environment previously cleaned using 10% bleach solution, rinsed with ethanol and ultrapure water. Each core of 10 cm diameter was sliced into five depth layers: 0–1, 1–3, 3–5, 5–10, and 10–15 cm. The slicing was introduced as, from coastal to abyssal settings where this was investigated, the depth in the upper centimetres of sediment has been underlined as a determinant parameter for the distribution of biodiversity across prokaryotes (Hoshino et al., 2020; Trouche et al., 2021), protists and fauna (Guardiola et al., 2016; Román et al., 2019), driven by organic-matter supply, redox zonation and bioturbation (Flach and Heip, 1996; Jorissen et al., 1995; Vanreusel et al., 1995). Slicing was performed using spatulas also bleached and rinsed with ultrapure water before each use. Horizons (slices of sediment) were transferred into zip-lock bags, homogenized, and frozen at -40°C on board before being shipped on dry ice to the laboratory where they were kept at -80°C. For the first layer, the overlying water of the core (approximatively 1cm height) was kept and homogenised with the 0-1 sediment layer. Field controls were prepared onboard: empty zip-lock bags were opened on site and submitted to the same condition as sample across the whole transport and laboratory procedures.

### 2.2. Environmental features and trawling pressure indices

Data describing the fishing effort, the impact of current, and the nature of the sediment was obtained following the steps detailed here below (data available in Supplementary Table 1)

#### 2.2.1. Swept Area Ratio (SAR)

The Swept Area Ratio (SAR) evaluates fishing pressure on the seabed. It is calculated from the interpolated trajectories of fishing vessels, derived from the Vessel Monitoring System (VMS) and the detection of fishing activities based on the vessel speed (Georges et al., 2025). These linear fishing actions are multiplied by the width of the fishing gear, leading to abrasion values at 1’x1’ (<4 km^2^) resolution integrated over a specific temporal duration. In the present study the SAR was determined for each station across three temporal integration periods from 2012 to 2021 (Table 1).

**Table 1.** Variables used to represent the fishing pressure indices and environmental descriptors at each station:

| Variable name | Description |
| --- | --- |
| **SAR_P90** | 90th percentile of the Swept Area Ratio over 10 years (P90 2012-2021 SAR) at the site, representing the long-term effect of trawl fishing |
| **SAR_wmean_2019_2021** | Weighted mean of the SAR at the site, over 3 years (2019-2021), representing the medium-term effect of trawl fishing |
| **SAR_072021** | SAR at the site, integrated over the month before the survey, representing the short-term effect of trawl fishing |
| **EP_h_per_month** | Fishing effort (number of hours of trawling) at the site, in the month preceding the survey |
| **EP_h_per_week** | Fishing effort (number of hours of trawling) at the site, in the week preceding the survey |
| **Dx_50** | Median of the distribution of sediment grain size |
| **Bedstress** | 90th percentile of shear stress on the seabed due to waves and currents |
| **Water_depth** | Depth of the station |

#### 2.2.2. Fishing effort (EP)

Another quantification of the trawl fishing intensity, the fishing effort (EP: “*effort de pêche*”), was also used. This data is obtained by the reconstruction of individual trawling fishing vessel trajectories from VMS, allowing the quantification of the fishing effort, within a buffer zone of a 50m radius around the station. This fishing effort is expressed as a number of hours of trawling on the site per month (over the 2019-2021 time period) or per week (over the months of July and august 2021).

#### 2.2.3. Particle size distribution

Particle size analyses were performed for each 1cm horizon of the core on bulk samples using a Malvern Mastersizer 3000 equipped with a wet dispersion unit (Hydro EV) (UMR LPG, Angers). The samples were first soaked in water and mechanically agitated for 30 seconds at 17 Hz. Laser obscuration was kept within the optimal range of 5–15%. Particle size distributions were calculated using the Fraunhofer approximation, assuming spherical and opaque particles (ISO 13320, 2020). For each sediment sample, three consecutive measurements were acquired. In accordance with the ISO 13320 (2020) standard, the repeatability of the measurements was validated by ensuring that the coefficient of variation (CV) remained below 3% for the median diameter (D50) and below 5% for D10 and D90. In exceptional cases, values of 5% and 10%, respectively, were accepted, in accordance with the protocols of studies on marine sediments conducted by (Blott and Pye, 2012; Sperazza et al., 2004). Key textural parameters, including volumetric percentiles (volume percentages of sand, silt, and clay; median particle size D50…), were computed using GRADISTAT v 9.1 software (Blott and Pye, 2001).

#### 2.2.4. Bedstress

The impact of natural forces (waves and currents) on the sea bed has been taken into account by calculating the seabed shear stress. The 3D MEDNOR hydrodynamic model (for current fields) and the MED01DEG model (for wave significant height, peak frequency, peak direction and bathymetry) were used to determine the hydrodynamics and to calculate the resulting shear stress on the sea bed, summarised at each station by the 90th percentile over the years 2001 and June 2007 to April 2009 (limited to available wave data periods) (Rivier, 2010).

### 2.3. Assessment of prokaryote, protist, and Animalia communities through eDNA

DNA was extracted using the Power Max Soil Kit (Qiagen) and 10g sediment per extraction. DNA was quantified using qPCR to decide on the dilution to apply for each marker (dilution by five for 18S-V4, 18S-V9 and 16S-V4V5, and dilution by two for 18S-V1 & COI). Controls were added by pouring 10ml MilliQ water into the first step tube of the kit, either directly from the manufacturer recipient (extraction control) or after having rinsed the internal sides of the empty bag prepared as a field control on the boat. Controls bags were filled with 10ml MilliQ water that was used for DNA extraction. Metabarcode libraries were then constructed using the protocol Metafast (one PCR followed by ligation of the tags). For all PCR, the PCR reaction was composed of 2 µl diluted DNA, 10 µl AmpliTaq Fold® 360 Master Mix’ (Fisher scientific), 0.16 µl BSA at 20 mg/L and 5.84 µl Water plus 2µl primers at 5µM each. For all batch of PCR reaction, a PCR control was added with the same mix as sample and 2 µl milliQ water instead of DNA. The PCR cycling was then specific to each marker. For the COI marker: a touchdown PCR with a pre-denaturation step at 95°C for 10 min, followed by 16 cycles of denaturation at 95°C for 30 s, annealing from 62°C to 47°C for 30 s, elongation at 72°C for 1.5 min, then 44 cycles of denaturation at 95°C for 30 s, annealing at 46°C for 30 s, elongation at 72°C for 1.5 min, and a final elongation at 72°C for 7 min. For the 18S-V4 marker: a touchdown PCR with a pre-denaturation step at 95°C for 10 min, followed by 10 cycles of denaturation at 95°C for 30 s, annealing at 53°C for 30s, elongation at 72°C for 1.5 min, then 30 cycles of denaturation at 95°C for 30 s, annealing at 48°C for 30 s, elongation at 72°C for 1.5 min and a final elongation at 72°C for 7 min. For the 18S-V1 a pre-denaturation step at 95°C for 10 min, followed by 40 cycles of denaturation at 95°C for 30 s, annealing at 45°C for 30 s, elongation at 72°C for 1.5 min and a final elongation at 72°C for 7 min. For the 18S-V9 a pre-denaturation step at 95°C for 10 min, followed by 32 cycles of denaturation at 95°C for 30 s, annealing at 57°C for 30 s, elongation at 72°C for 1 min and a final elongation at 72°C for 7 min. For the 16S-V4 a pre-denaturation step at 95°C for 10 min, followed by 40 cycles of denaturation at 95°C for 30 s, annealing at 53°C for 30 s, elongation at 72°C for 1.5 min and a final elongation at 72°C for 7 min.

Ligation and sequencing were then performed at Fasteris on a Illumina Novaseq.

### 2.4. Bioinformatic analysis

Bioinformatic analysis were performed on Ifremer cluster (Datarmor) using an update of the published Unix shell script (Brandt et al., 2021) available at (https://gitlab.ifremer.fr/abyss-project/abyss-pipeline), details of versions and sets of parameters are provided in Supplementary Table 2.

Briefly, after using cutadapt to remove adapters and trim sequences, the analysis started with a denoising step performed using DADA2 (Callahan, et al. 2016) to produce an Amplicon Single variants (ASVs) table. Based on negative controls at all steps (sampling blanks, extraction blanks, PCR blanks), a R script was used on ASVs or OTUs tables to process to a decontamination of the dataset using the package decontam (Davis et al., 2018), and a correction for tag switching (Wangensteen et al., 2018). For COI and 18S markers, ASVs are then clustered using Swarm v3 algorithm (Mahé et al., 2021) to aggregate sequences iteratively depending on their number of nucleotide differences. Considering the different resolution power of the markers used, 18S markers being less discriminant than COI (Tang et al., 2012), we used d=1 and the *fastidious* option of swarm for all 18S markers, enabling to account for possible ghosts ASVs and to aggregate ASVs distant of more than d, and adopted d=6 for COI to produce a table of operational Taxonomic units (OTUs). A post-clustering curation was then applied to COI data using MUMU ((Mahé, n.d.), a new implementation of LULU (Frøslev et al., 2017)) to minimize the bias due to the existence of paralogs and pseudogenes or the persistence of sequencing errors, setting a minimum occurrence of 90% and a minimum similarity threshold of 84%.

Taxonomic assignments of ASVs and OTUs were performed with BLAST+ using relaxed thresholds (BLAST+ v2.6.0 based on minimum similarity 70 and coverage 80) on the COI marker, against the MIDORI GB256 (Machida et al., 2017). The CREST4 v4.4.2 (Lanzén et al., 2012) was used with the implemented silvamod138pr2 database (based on Silva138 for SSUr RNA marker genes (Quast et al., 2012), using the blast algorithm (blastn as implemented in CREST4) for all 18S markers, and the vsearch algorithm (usearch_global parameter as implemented in CREST4) for 16S. For protist identification, the 18S-V9 sequences were also reassigned with RDP classifier implemented in DADA2 on the PR2 version 5.1.0 database (Vaulot and González-Miguéns, 2025), with a taxonomic assignment workflow (https://gitlab.com/metabarcoding_utils/taxonomic-assignment), keeping only assignations with a confidence higher than 60. Gaps in the expected 7 ranks COI taxonomic assignment from the pipeline were completed with the help of the WoRMS database ((WoRMS Editorial Board, 2026), accessed on January 2026 with the package worrms (Chamberlain and Vanhoorne., 2017)), and using a custom R script.

The 4 technical PCR replicates per sample were then merged, cells (ASV x sample) with less than 5 reads were set to zero, and samples totalling less than 9000 reads for 18S-V9 and 16S-V4V5, and 5000 reads for COI, 18S-V1 and 18S-V4 were removed.

For the study of benthic prokaryotes (Bacteria and Archaea), the 16SV4V5 marker was used and the ASV table was filtered to keep only those assigned to the phylum level, and belonging to the bacteria or archaea kingdoms.

For protist the 18S-V9 marker was used and ASV were filtered based on their class, according to a list of protist class extracted from the pr2 database, in which Streptophyta and Metazoa were filtered out.

For Animalia, the COI, 18S-V1, 18S-V4 and 18S-V9 markers were jointly used for their complementarity among phylum. First, only OTU assigned to an Animalia phylum were kept. Chordates were then split into Vertebrata and Tunicata due to the important ecological differences between these two subphyla. The WoRMS database was used to filter the OTU from each marker and keep only marine Animalia removing terrestrial organisms in the following classes: Mammalia, Insecta, Arachnida and Aves. Then, for each Animalia phylum, the marker that identified the largest number of OTU was retained, allowing the combination of the information from different markers and the OTU table was set up to presence absence.

Finally, for Animalia and protists, the taxonomic assignation of ASV/OTU was cross referenced with the WoRMS database in order to characterise them as benthic or pelagic (WoRMS attributes, functional group), or unknown when no information was available.

### 2.5. Statistical analysis

#### 2.5.1. Diversity indices

All subsequent statistical analyses were done in R v4.5.0, using the phyloseq v1.52.0, (McMurdie and Holmes, 2013) and vegan v2.7-1, (Oksanen et al., 2001) packages. Visuals were produced with ggplot2 v3.5.2, (Wickham, 2016) and gt v1.3.0, (Iannone et al., 2020). Environmental parameters considered for each sample were sediment horizon (depth in the sediment core), trawling disturbance on the seabed (SAR and EP at the station), the natural disturbance on the seabed (bedstress at the station) and the sediment grain size (Dx_50, the median size of particles) for each layer of sediment.

Alpha diversity was computed with the vegan package in R, using the Shannon index for prokaryotes and protists, and the richness as the observed number of OTU for Animalia. The Pielou index representing eveness in prokaryote and protists communities was computed the same way.

Beta diversity of the communities was illustrated by performing a PCoA on the samples (with the triplicates of cores merged together, by summing reads per station and per horizon) with the wcmdscale function from the vegan package. For Animalia, this PCoA was based on the Jaccard distance matrix, based on the number of OTU per sample. The distance matrix computed with the vegdist function, after having discarded the OTU present in only one sample. For protists and prokaryotes, the read numbers were first normalised using the metagenomeSeq package (Joseph Nathaniel Paulson, 2017; Paulson et al., 2013), a Bray-Curtis dissimilarity matrix was calculated on the normalised reads with the distance function from the phyloseq package, and the PCoA was computed.

#### 2.5.2 Environment-Fishing indices relationships

The influence of standardized environmental and fishing descriptors (scaled to mean=0 and standard deviation=1) on the median particle size of the sediments was evaluated using glmmTMB v1.1.13, (Brooks et al., 2017) with the following formula, under a gaussian distribution, since the median particle size is a continuous numerical variable:

D*x*_50 ∼ *horizons* + *bedstress* + *water*_*depth* + *SAR*_072021 + *SAR*_*P*90 + *SAR*_*wmean*_2019_2021 + *EP*_*h*_*per*_*month* + *EP*_*h*_*per*_*week* + (1 | *station*)

A mixed model was chosen to account for the grouping effect of the station on the median particle size

#### 2.5.3 Environment-Fishing indices-diversity relationships

The correlation between the environmental and fishing variables, and the alpha diversity of the 3 taxonomic groups (prokaryote, protist, Animalia) were evaluated using generalised linear mixed models (glmm), on standardized environmental and fishing descriptors (scaled to mean=0 and standard deviation=1, to put them on the same scale and enable easier comparison of their association with the alpha diversity of the communities). The use of *glmm* allows accounting for the random effect of the station on the alpha diversity of the samples, and treating data that does not follow a normal distribution (e.g. the number of OTU in Animalia).

Models evaluating the link among the alpha diversity of the three taxonomic groups and environmental and fishing descriptors were computed. For these models, the grouping of the samples by station, and also by cores within the stations, was represented by a nested random effect of stations, and cores within stations (random effect: (1 | station/core)), in *glmm*. However, while this random effect made sense according to the sampling structure, it rendered the models singular. Therefore, models were computed without a random effect: generalised linear models (*glm*) for the counts of OTU in Animalia, that require a negative binomial distribution (discrete and positive data, with overdispersion), or simple linear models (*lm*) for the Shannon index computed for prokaryote and protist data, that require a gaussian distribution (continuous numerical values). To better fit a gaussian distribution, the Shannon index of the protist data was transformed: inverse transformation (1 /x) on the reflection of the distribution (max(shannon)-x). To compute the models, the glm.nb function from the MASS package (7.3-65) was used for Animalia, and the lm function from R stat for protists and prokaryotes.

The relationship between alpha diversity in Animalia, and alpha diversity in protists and prokaryotes was examined with the following model (under a negative binomial distribution):

*alpha_observed_Animalia* ∼ *Shannon_protist* + *Pielou_J_protist* + *Shannon_prokaryote* + *Pielou_J_prokaryote*

Similarly, the link between the alpha diversity in protists and prokaryotes was modelled under a Gaussian distribution (with transformation of the protist Shannon index), with the following formula:

*Shannon*_*protist* ∼ *Shannon*_*prokaryote* + *Pielou*_*J*_*prokaryote*

Then the effect of all the environmental variables on the alpha diversity of organisms (i.e. the number of OTU for Animalia, and the Shannon index for protist and prokaryotes) was evaluated, through a single model, computed separately on each of the three taxonomic groups.

*alpha*_*diversity* ∼ *bedstress* + *horizons* + *SAR*_072021 +*SAR*_*P*90 + *SAR*_*wmean*_2019_2021 + *EP*_*h*_*per*_*month* + *EP*_*h*_*per*_*week*

For protists and Animalia, for which ASV and OTU could be classified as representing benthic or pelagic species, the models were computed separately on the three compartments: on all ASV/OTU, on ASV/OTU identified as benthic, or on ASV/OTU identified as pelagic.

To identify the environmental variables most strongly associated with the alpha diversity of the different organisms, models based on all possible combination of the explanatory variables were fitted and compared. Candidate models were ranked using the corrected Akaike information criterion (AICc) which evaluates the trade-off between goodness-of-fit and model complexity (with a correction for small samples, compared to simple AIC). Model averaging was performed on the best supported models (delta AICc <=2 compared to the best model with the lowest AICc), and the averaged estimates of explanatory variables were retained as the representation of the most decisive environmental factors influencing alpha diversity. Model selection and model averaging were conducted using the dredge function from the MuMIn R package v 1.48.11 (Bartoń, 2010).

Model assumptions were assessed using the DHARMa (v0.4.7) (Hartig F, 2026) and performance (v0.16.0) (Lüdecke et al., 2021) packages. Simulated residuals were inspected visually and tested for uniformity, dispersion, and outliers. Multicollinearity among predictors was evaluated using variance inflation factors (VIFs). Model performance was quantified using R^2^ values (or pseudoR^2^ from Nagelkerke for the glm, calculated with the performance package).

The link between the environmental variables and the communities was also tested by distance-based redundancy analysis (dbRDA, dbrda function from vegan) on the distance matrices. A forward selection of predictors was operated on the rdRDA based on R^2^ with the ordiR2step function from vegan. Variance partitioning was conducted on the dbRDA, to distinguish the effect of trawling (accounting for SAR_P90, SAR_wmean_2019_2021, SAR_072021, EP_h_per_month and EP_h_per_week) from the effect of natural mechanical constraint on the seabed (bedstress) on the community structure.

### 2.6. Cross referencing between metabarcoding data and data from the literature

Names of organisms previously morphologically identified in studies of the benthic environment in the Gulf of Lion (references used available in Supplementary Table 3), and by morphological surveys conducted of benthic mega-epifauna, macrofauna and meiofauna for the IMPEC project (Barras et al., 2025) were extracted. Taxonomic identifications from these sources were heterogenous, ranging from species-level identifications to broader taxonomic groups, in some cases as high as the phylum level (e.g., Nemertea). For each taxon identified through the literature and morphological surveys, the complex taxonomic hierarchy was retrieved from the WoRMS database. This way, the correspondence with the taxonomic assignments of OTUs could be done. This correspondence used the most precise taxonomic level available for organisms mentioned in literature or in the morphologically survey, except for species-level identification. In this case, the comparison started at the genus level because broad metabarcoding species assignation can be unreliable at the species level. A taxonomic tolerance margin was implemented in the comparison, to account for differences in taxonomic resolution between morphological identification and eDNA assignments. When no correspondence was found at the initial taxonomic level of the morphological identification, the comparison was extended to the upper taxonomic level. For instance, if an organism was identified morphologically to the genus level, but no OTU was assigned to that genus, the OTU table was subsequently screened for assignments to the corresponding family.

## 3. Results

Details on the read numbers and ASV/OTU numbers retained along the bioinformatic pipeline is available in Supplementary Table 4.

For Bacteria and Archaea, the 16S marker outlined a total of 201 706 ASVs, mostly identified at the order level (Supplementary Figure 1), and belonging to 106 different phyla. For protists, a total of 10 306 ASVs were retained with 18S-V9, mostly identified at the species level (Supplementary Figure 2), and belonging to 37 distinct Subdivisions. Among these protist ASV, 2 681 could be identified as benthic ASVs, and 5 092 as pelagic ASVs, based on cross referencing of the taxonomic assignment with the WoRMS database. For Animalia, a total of 6 342 OTU was obtained with the combination of the four markers (COI, 18S-V1, 18S-V4 and 18S-V9), mostly identified at the species level (Supplementary Figure 3). Among these Animalia OTUs, 5 090 could be identified as benthic, and 378 as pelagic, based on cross referencing of the taxonomic assignment with the WoRMS database. For the semi quantitative datasets (prokaryotes and protists), the rarefactions curves confirmed that sequencing and sampling efforts captured most of the sample diversity (Supplementary Figure 4).

### 3.1. Composition of communities in the 8 stations from the Gulf of Lion

First, the analysis of environment variables showed that stations A and C belong to a “Mediterranean circalittoral coarse sediment” habitat (MC35), whereas stations E, G, J belong to the “Mediterranean offshore circalittoral mixed sediment” habitat (MD45) and stations K, O, P belong to the “Mediterranean offshore circalittoral mud” habitat (MD65), according to the EUNIS habitat classification. heoretical prediction of the MD451 habitat used to plan the sampling supposed that stations A, C, E, G, J should correspond to the MC35 habitat but the span of this habitat was underestimated near the stations A and C (Barras et al., 2025). In terms of the available environmental data, the main difference of station A and C compared to the other stations lies in the higher values of bedstress and coarser sediment grain size (seen in Dx_50 values), that can be linked to the location of these stations close to a steep slope (Figure 1).

In fact, the stations A and C display differences in terms of alpha and beta diversity of communities, while the 6 other stations exhibit very similar profiles (Figure 2, Supplementary Figure 5).

**Figure 2:**
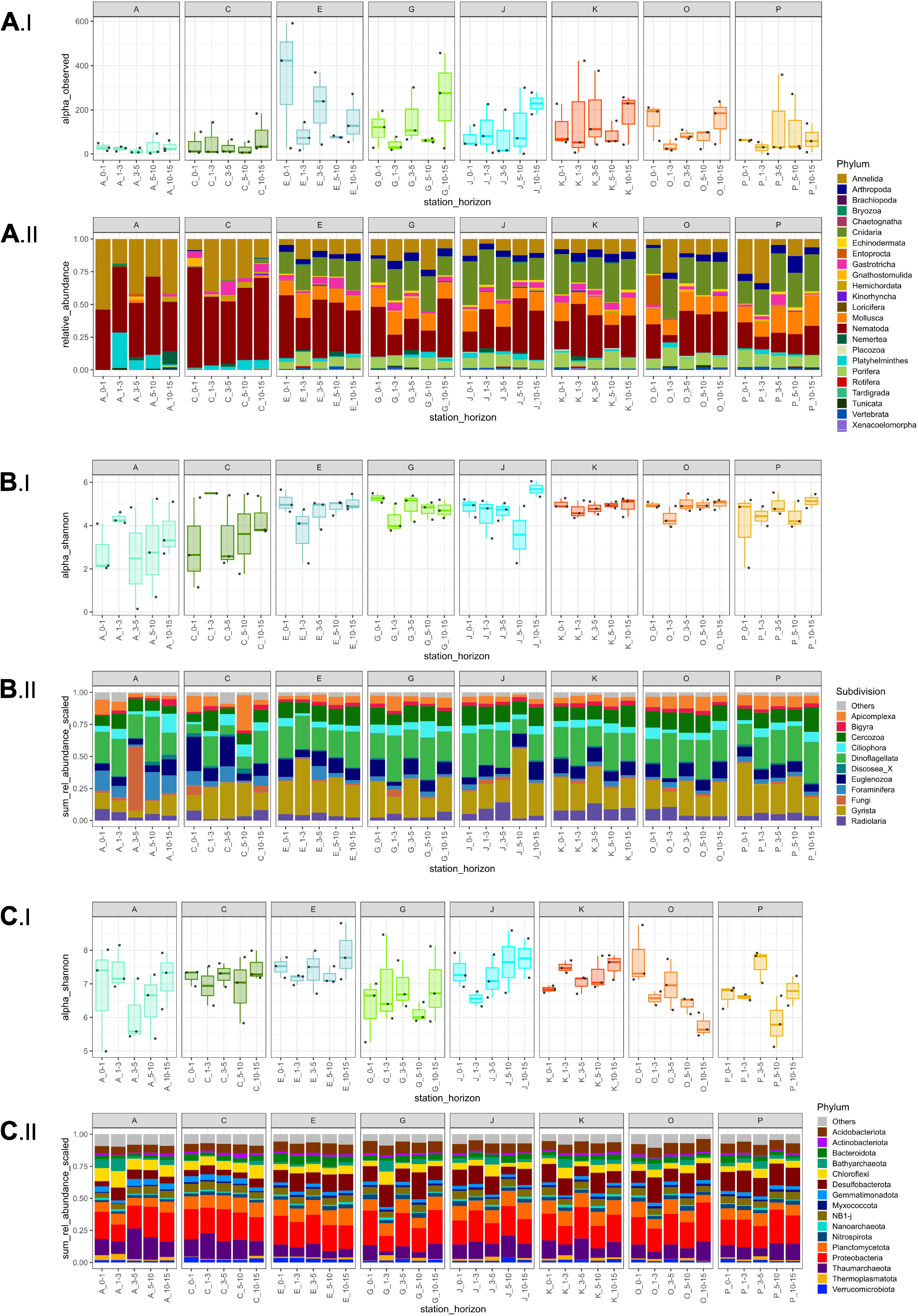
Overview of the sediment layers community content in all stations, for Animalia (A), protists (B) and prokaryotes (C). The alpha diversity (I) is represented by the Shannon index for prokaryotes and protists, and by the number of OTU (observed alpha diversity) for Animalia, ordered by increasing layer depth in the sediment. The barplots of taxonomic abundance (II) display the relative abundances in read numbers (prokaryotes, protists) or OTU numbers (Animalia) for each major Phylum for prokaryotes and Animalia, or Subdivision for protists. For protists and Animalia, the benthic or pelagic lifestyle of taxonomic groups was extracted from the WoRMS database (Supplementary Figure 6). For Animalia, the alpha diversity boxplot and the barplot of Phylum composition are based only on Animalia OTU identified as benthic in WoRMS, whereas for protists and prokaryotes, the plots are based on all the ASV.

In terms of alpha diversity, Animalia are the taxonomic group displaying the starkest difference between stations A and C and the other stations (Figure 2 A). Station A has a significantly lower alpha diversity than stations E, G and O (significant Kruskall Wallis test over all stations, with p-value of 1.76e-4, and post hoc Tukey to test pairwise differences: A-E p-value of 0.009, A-G p-value of 0.019 and A-O p-value of 0.026). It also has a different taxonomic composition, with almost no Arthropoda, Cnidaria, Mollusca and Vertebrata (respectively 0.29%, 0%, 0.24% and 0.24% of the OTUs) that are quite common in other stations, and has a greater proportion of the OTUs being assigned to Annelida (37% of the OTUs) or Nematoda (48% of the OTUs) phyla. Station C displays a similar, yet more nuanced profile, with slightly higher proportion of Cnidaria and Mollusca (respectively 1.19% and 0.89% of the OTUs), yet this station is dominated by Nematodes (60% of the OTUs). This difference between stations A and C and the other stations is also apparent on the PCoA of Animalia beta diversity across stations and sediment horizons (Supplementary Figure 5 A.I).

This difference between stations A and C and the others is also suggested on the PCoA ordination and the plot of alpha diversity of protists, but is non-significant in the latter (Kruskall Wallis test p-value of 0.06) (Figure 2B, Supplementary Figure 5 B.I). For prokaryotes, no clear significant difference of diversity between stations is detected (ANOVA over alpha diversity in stations is significant with a p-value of 0.0128, but the comparisons of Tukey’s test are not), yet, a separation between stations A and C and the other stations can be observed on the PCoA describing beta diversity (Supplementary Figure 5 C.I).

Along the depth of the sediment, we see no clear difference of alpha diversity in the communities (p-value of the Kruskall Wallis test between horizons: for Animalia: 0.11, for protists: 0.29, for prokaryotes: 0.31), and no clear difference of phylum (or subdivision) composition (Figure 2), or position of the horizons on the PCoA either (Supplementary Figure 5 A.II, B.II, C.II). Interestingly, there is also no visible trend in terms of the ratio between benthic ASV vs pelagic ASV identified for Animalia and protist in the sediment, while it could be expected that layers closer to the water column could contain more pelagic DNA (Laroche et al., 2020) (Supplementary Figure 6). Moreover, particle-size distributions varied between sediment horizons, but for most cores, no consistent trend in particle-size characteristics was observed with increasing sediment depth (the layers ordered by median particle size value are not necessarily following an ascending or descending order of depth, Figure 1). These elements suggest a homogenisation of the sediment layers in these eight stations, but it is hard to state if this homogenisation is mostly due to the trawling activity or to hydrodynamic forces.

### 3.2. Interactions between the three taxonomic groups

A significant association was found by glm only for the protist community metrics: with Animalia OTU richness which was positively and significantly associated with the protist Shannon index, and negatively and significantly associated with protist evenness (Pielou index) (Supplementary Table 5). A model including only the protist Shannon index and evenness as predictors accounted for a large proportion of the variance in Animalia OTU richness, as indicates the pseudo-R^2^ of 0.87 from the model only with protist predictor variables. Prokaryotes alpha diversity and evenness, by contrast, showed no significant association with Animalia diversity, and accounted for only a small proportion of the variance in the protist Shannon index (only the Shannon index of the prokaryotes is significantly associated to the Shannon of the protists, in a model with a R^2^ of 0.03) (Supplementary Table 5).

### 3.3. Effect of the environmental variables on the Animalia, protist and prokaryote communities

Seabed trawling alters the sediment structure by homogenisation of sediment layers, flattening of seabed relief, bed armouring and resuspension of large volumes of sediment (Martín et al., 2014b; Schwinghamer et al., 1998). This adds to the direct mechanical effect of trawling on benthic communities, an indirect effect mediated by the reworking of the sedimentary environment. While the direct mechanical effect is acting mainly on large-size organisms, the physical and biochemical perturbations in the sediment can also affect benthic microorganism communities. Therefore, among the different environmental variables available, the ones describing the trawling and the hydrodynamic disturbance of the seabed can have an impact on the structure of the sediment and therefore covary with the granulometry, which can in turn impact the benthic communities. This relationship was tested with a linear mixed model. Almost all variance in sediment D50 across samples was accounted for by the other variables (marginal R^2^=0.88), with bedstress alone accounting for 36% of the explained variance, and all trawling variables accounting together for 41% of the explained variance (hierarchical partitioning analysis, see Supplementary Table 5). Grain size information was therefore excluded from subsequent models, as most of its relationships with the communities is captured indirectly through the other environmental variables. This way, the modelled relationship between trawling pressure variables and the benthos will account for both the direct effects of trawling and its indirect effects mediated by changes in sediment characteristics.

The remaining environmental variables, describing natural (bedstress) or trawling-related (SAR and EP) seabed constraints were tested against the alpha diversity indices of the three taxonomic groups (biplots of the trends between these variables and the alpha diversity available in Supplementary Figure 7), using (generalised) linear models, with separate models for benthic and pelagic ASV/OTU where available for protists and Animalia. The first model computed for each taxonomic group (and benthic/pelagic category if available) was a model with all the different predictors. Predictors were selected by AICc-based model selection and averaging (models within ΔAICc < 2) to retain only the most influential ones. SAR_P90 and SAR_wmean_2019_2021 showed moderate collinearity in the full models; both were retained to capture different timescales of trawling pressure, and results involving them should be read with this in mind. Retained predictors, their significance, and their direction of association with each alpha diversity metric are reported in Figure 3 and summarized in Figure 4. For Animalia, the model on benthic OTU only is the one retained, while for protists, the model on all ASV is retained, since many ASV could not be assigned to the benthic or pelagic compartment when cross-referencing them with the WoRMS database (Supplementary Figure 6).

**Figure 3:**
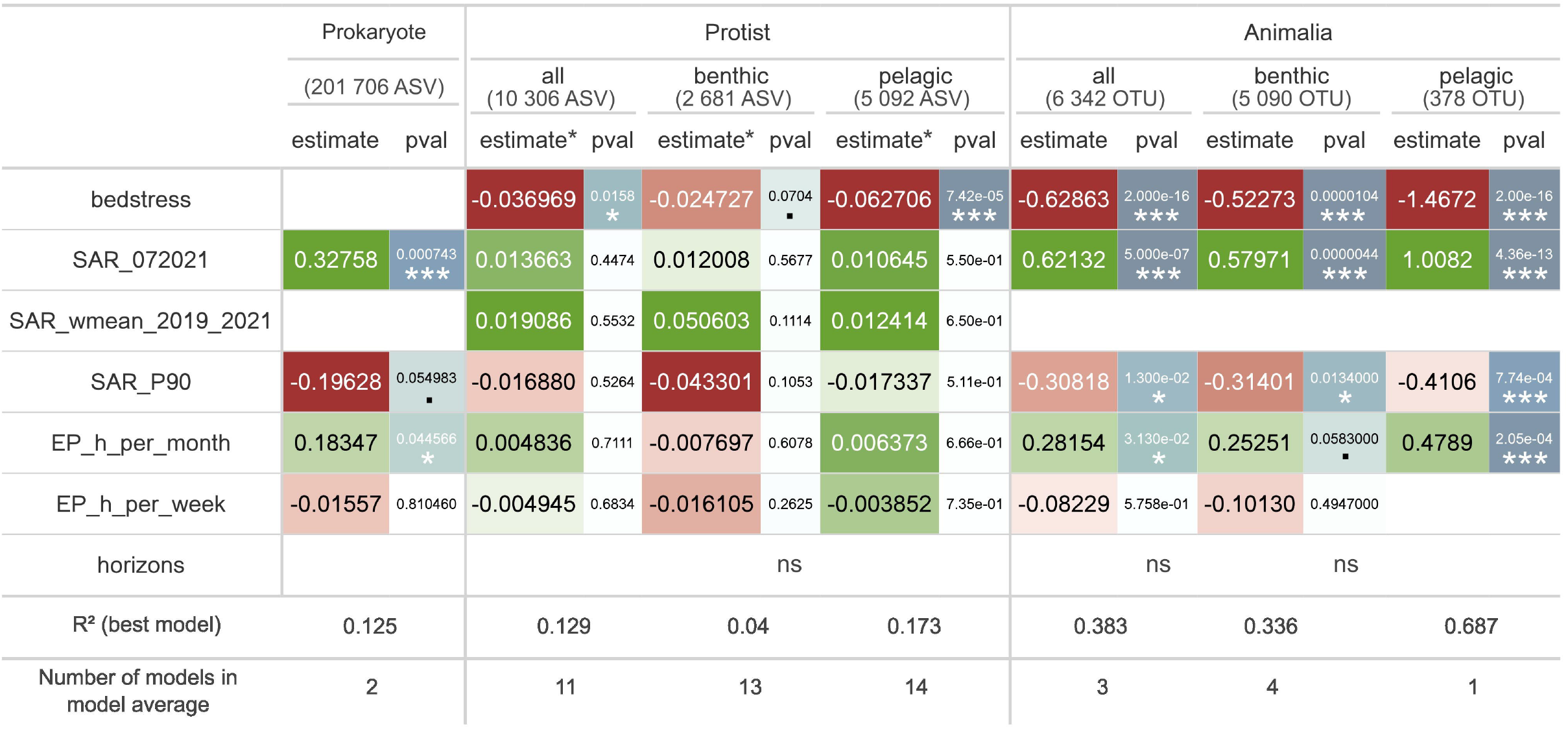
Physical disturbances significantly associated with the variation of alpha diversity in the three types of organisms, from model averaging based on the best subset models according to the AICc selection on the GLMM with all predictive variables. For protists and Animalia, the models based on all taxonomic units (ASV and OTU, respectively), are displayed, as well as models based only on benthic or pelagic groups. For each model the estimated regression coefficients of the model on each variable (the estimates) are specified, accompanied by their associated p-value. For protists, the “estimates*” indicates that the models were computed on a transformed Shannon index. The estimated coefficients are based on scaled values of the explanatory variables, and are coloured depending on their values (green: positive, red: negative). P-values are displayed for each predictive variable retained in the final model, with a darker colour for smaller p-values, and white for non-significant p-values. The abiotic variables considered are the following: bedstress: effect of the current and waves on the seabed; SAR= abrasion values integrated over different time period before the study: SAR_072021: in the month, SAR_wmean_2019_2021: in the three year or SAR_P90: in the past ten years; EP=effort de pêche (fishing effort), in the month or in the week preceding the study; horizons: depth of the sediment, cut in standardised layers: 0-1,1-3,3-5,5-10 and 10-15. When “ns” is specified for p-value of horizons, it means that the different horizons of the sediment are included in the averaged model, but none of them alone is associated to a significant value. The R^2^ (i.e. the variance of the alpha diversity that can be explained by the predictive variables selected, or the pseudo R^2^ for Animalia GLMs) specified in this table is the one of the best model. The number of models averaged to get to these estimates and p-values is also specified.

**Figure 4:**
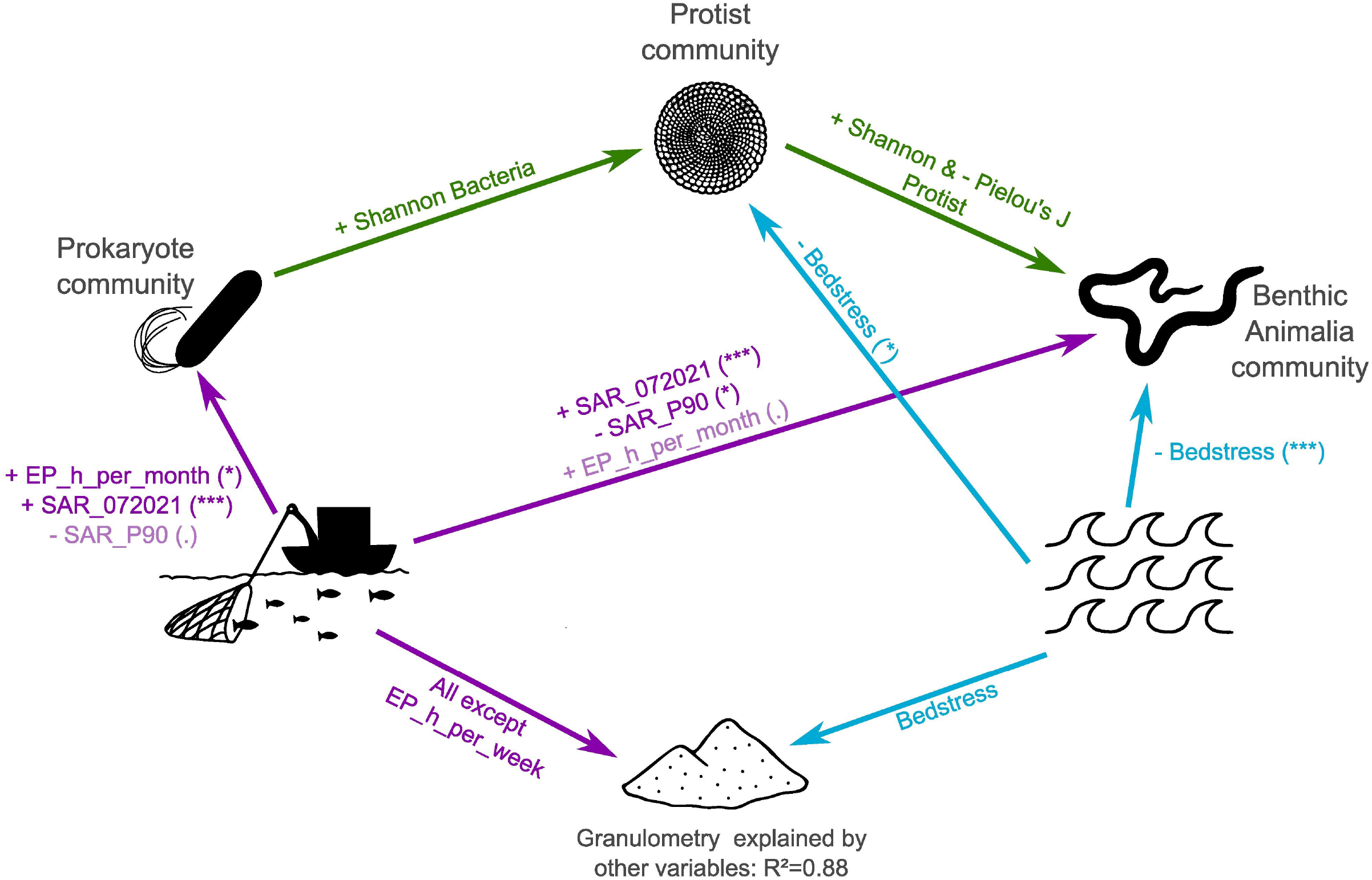
Schematic summary of the trawling and natural (bedstress) variables significantly influencing the alpha diversity of the different types of organisms. Variables listed on this figure were present in the averaging of the best models, according to the AICc, and significantly associated with alpha diversity variation (except for two predictors that were almost significant: SAR_P90 for prokaryotes: p-value of 0.054 and EP_h_per_month in Animalia: p-value of 0.058). The sign before the variables on the arrows shows the sign of the estimated effect of the variable (positive or negative impact on the diversity). For Animalia, the model on the alpha diversity reduced to benthic taxa was used, whereas for prokaryotes and protists, all ASVs were considered.

No significant association was found between protist alpha diversity and any trawling metric. For prokaryotes and Animalia, trawling metrics showed contrasting association with alpha diversity depending on the time scale. Prokaryotes alpha diversity was significantly and positively associated with short-term trawling: the more the area is trawled in the past month (SAR_072021 and EP_h_per_month), the more alpha diversity is observed for Bacteria and Archaea. Long-term trawling over the past 10 years (SAR_P90) was associated with lower alpha diversity of prokaryotes but this trend did not reach significance (p-value 0.054). A comparable pattern is observed for Animalia (whether benthic, pelagic or all datasets): alpha diversity was positively associated with short-term trawling (SAR_072021 (p-value 4.4e-6), nearly significantly positively associated with EP_h_per_month (p-value 0.058)), and significantly negatively associated with long-term trawling pressure (SAR_P90, p-value 0.013).

The hydrodynamic constraints applied on the seabed, the bedstress, was significantly and negatively associated with the alpha diversity of protists and Animalia respectively. This echoes to the difference in alpha diversity and Phylum composition observed between stations with higher bedstress constraint (A and C) and stations with lower bedstress, a pattern more evident for Animalia and protist communities. According to the analysis of variance partition analysis on the structure of the communities, bedstress has as much effect as all the trawling metrics combined on Animalia and protist communities, and more effect than trawling on prokaryote communities (Supplementary Figure 5).

Sediment horizon was occasionally retained in the top models, but was never significantly associated with alpha diversity after model averaging, indicating no detectable gradient in alpha diversity across sediment layers.

The distance-based redundancy analysis (dbRDA) showed that bedstress and SAR_072021 were the predictors retained by forward selection as significantly associated with community structure for all three taxonomic compartments. However, most of the between-sample variation in community composition remains unexplained, as shown in the variance partition analysis (Supplementary Figure 5 A.III B.III C.III).

## 4. Discussion

### 4.1. Complementarity between the metabarcoding approach and traditional morphological identification to evaluate benthic communities

The sediment-based eDNA metarbacoding on 119 samples from 8 stations on the continental shelf of the Gulf of Lion, combining 5 different metabarcoding markers, allowed characterizing 6,342 Animalia OTUs, 10,306 protist ASVs and 201,706 Bacteria ASVs.

The eDNA analysis enabled the identification of species listed as typical from the Gulf of Lion. The genera *Leptometra, Ophiura, Antalis* and *Haploops*, typical of the biocenosis of open-sea detritic bottoms (La Rivière et al., 2021) were detected by eDNA, in multiple stations (C, E, K for *Leptometra*, K for *Ophiura*, E for *Antalis*, E and K for *Haploops*). Three genera out of the four described as typical indicators of different benthic communities from the Gulf of Lion (Guille, 1970; Labrune et al., 2007), were also identified: *Spisula* (only P station), *Nepthys* (E, J, K, O stations) and *Nucula* (E, G, J, K stations). In the same way, many Polychaetes taxa described in studies of shallow soft bottoms of the Gulf of Lion (Bonifácio et al., 2019; Labrune et al., 2007; Rosenberg et al., 2003) were identified here, notably from the Lumbrineridae, Spionidae, or Oweniidae families.

In the context of the IMPEC project, morphological identification of mega and macrofauna taxa was also carried out at the same time as the benthic eDNA sampling, in stations A, C, E, G, J (Barras et al., 2025). Two distinct methods were used: processing of sediment samples for macrofauna morphological identification, and analysis of video transects for identification of megafauna located at the surface of the sediment. Occurrences of taxa found in all stations were compared with occurrences of the same taxon found by metabarcoding in each station, to determine to what extent the taxa targeted by morphological identification were also detected by the broad-range eDNA survey. It appears that the metabarcoding approach covers 61% of the megafauna taxa detected by video and 52% of the macrofauna taxa detected by sediment analysis (knowing that the taxonomic level of taxa considered is quite heterogenous: some taxa identified by morphological analyses can be Phylum, like Nematoda, while others can be at the Genus level, such as *Eunice*). The morphological data focuses mainly on Mollusca and Annelida; therefore, the morphological description of the benthic Animalia community is less exhaustive than the one obtained by metabarcoding, but also less precise, since for some taxa the morphological description only goes up the phylum level. Among the 24 benthic phyla detected by eDNA, 14 of them, such as Entoprocta, Gastrotricha or Gnatostomulida are not mentioned in the morphological analyses either because they are not searched for, notably because many meiofauna or microscopic taxa are extremely hard to detect and identify, or because they are simply undetected.

The identification of taxa by the different methods (morphological vs eDNA) was scrutinised station per station. Some taxa, present in all stations, are well identified by both methods. These taxa are mostly Polychaetes for the macrofauna identified by morphological sediment analysis, and Anthozoa for the video analysis of megafauna. The eDNA analysis also made it possible to detect some of the taxa at more stations than by morphological identification: for instance, Gastropoda, *Ophelina*, and *Chaetozone* were detected at only one or two stations by morphological survey of the sediment, but at all stations with the eDNA analysis. On the other hand, for some taxa, such as *Leptometra* that are typical of Mediterranean open-sea detritic bottoms, the morphological video survey enabled their identification in 4 stations (A, C, E and G), while their eDNA signature was only found in two of these stations (stations C and E).

Overall, eDNA and morphological analyses appear again quite complementary (Cowart et al., 2015; Zhang et al., 2025), with the morphological analysis providing detailed information on some phyla of interest, while the eDNA analysis provides a broader view of the diversity of benthic Animalia at these sites. Despite the incomplete overlap between the two approaches, they show similar trends in terms of the community structure in the five stations they share in common. Indeed, the morphological analyses in various taxonomic groups (e.g. Nematoda, Macrofauna) show a clear separation of the communities between stations A, C and E, G, J, but does not find that any of the trawling metrics significantly explain the community structure, once other environmental factors are taken into account (Barras et al., 2025).

### 4.2. Similar impacts of trawling across the Tree of Life, with opposite time scale effects

Cross-referencing the eDNA dataset with trawling activity metrics revealed significant statistical associations with trawling pressure for two of the three taxonomic groups studied. Prokaryotes and Animalia — despite their marked phylogenetic, physiological, and life-history differences — displayed a strikingly similar pattern of response to trawling, both showing contrasting responses depending on the temporal scale of trawling pressure considered.

First, against our initial expectations, especially for larger and longer-lived Animalia that are more sensitive to trawling perturbation (Jennings et al., 2001; Kaiser et al., 2006; Queirós et al., 2006), short-term trawling (i.e. during the past month) is associated with higher alpha diversity in both prokaryotes and Animalia, as recently reported for prokaryotes and protists (yet not animalia) in deeply trawled area of the North Sea (Seidel et al., 2026). Communities can be modified by the side effects of trawling, without any loss of alpha diversity, through several mechanisms that are not mutually exclusive. First, the passage of the trawl generates a resuspension of nutrients buried in the sediment, an oxygenation of deeper layers in the sediments and an increase in carbon cycling, which can boost the activity of communities and explain the increase in diversity (Bradshaw et al., 2024; Jennings et al., 2001; Morys et al., 2021; Polymenakou et al., 2005; Watling and Norse, 1998). Second, the attraction of scavengers on the site (Kaiser and Spencer, 1996; Tillin et al., 2006) or the release of competition and predation, due to the damage done to some organisms (Jennings et al., 2001) can lead to a short-term increase in alpha diversity. Finally, the homogenisation of the sediment layers by the trawling disturbance can artificially inflate the overall diversity in each layer, by pooling together species initially inhabiting different layers.

The last hypothesis is supported by the lack of consistent vertical layering of biodiversity levels in our data, as well as the consistent proportions of pelagic protists and Animalia found across all sediment depths, hinting at a mixing of the organisms, or of their DNA itself, in the sediment layers. This is at odds with what is usually reported for undisturbed benthic sediments: a decrease of the diversity of organisms in deeper layers (Flach and Heip, 1996; Good et al., 2022; Guardiola et al., 2016), and a vertical layering of communities, notably along the redox gradient for microorganisms (Jorgensen et al., 2012). Such homogeneous distribution of the sediments may be expected as a result of the repetitive trawling disturbance in the area, which has been shown to affect vertical stratification of sediment biogeochemical properties (Martín et al., 2014a; Pusceddu et al., 2014). However, the lack of a control area free from trawling prevents us from drawing firm conclusions regarding the relative contribution of trawling and other disruptors of the sediment layering, such as bioturbation or hydrodynamic forces. This calls for further studies with a special effort put into the gathering of controls, even in extensively swept areas.

Long-term trawling pressures, on the other hand, appear to have a deleterious effect on the diversity of organisms on both prokaryotes and Animalia, with more robust statistical support for Animalia. As mentioned previously, most Animalia are more sensitive than protists or prokaryotes to the repeated disturbance caused by trawling (Bradshaw et al., 2024). Indeed, organisms of small size have a greater chance of being displaced rather than damaged or killed, and their short generation time allows them to rapidly recolonise the seabed after the disturbance (Jennings et al., 2001; Schratzberger and Jennings, 2002). This effect on Animalia is in line with previous findings suggesting that bottom-trawling pressure selects for species with faster life history and smaller body size (Jennings et al., 2001; Queirós et al., 2006). However, it is noticeable that prokaryote alpha diversity as well, despite short generation size and body size, displays a nearly significant negative association with long-term trawling intensity. This suggests that bottom-trawling might slowly erode microbial diversity (Bonthond et al., 2023), even though no consensual effect of trawling on bacterial communities has yet been determined, with some studies even finding no significant effect of trawling on bacterial communities (Bradshaw et al., 2024).

Contrastingly, no trend was observed in protist alpha diversity. This may reflect either the lack of effect of bottom trawling on the eDNA signature of this compartment, possibly due to the strong influence of pelagic (and thus, in this case, sinking) organisms, or a statistical limitation resulting from the transformation required to fit the models to the protist data, that might have reduced our power to detect a linear association with the environmental variables.

### 4.3. Pervasive trawling effects across different environmental contexts

The hydrodynamic constraints on the seabed also show an effect on the diversity of the benthic communities and seem to be size dependent, since Animalia are the most impacted. This influence of the bedstress on the alpha diversity of organisms is particularly emphasized in this study by the stark ecological singularity of stations A and C, which are highly disturbed by hydrodynamic currents. The observed correlations between bedstress and the diversity and structure of the communities underline bedstress as a shaping force of benthic communities both through direct (a mechanical constraint) and indirect effects (on granulometry, the washing out of nutrients…) (Thrush et al., 2021). But this correlation could also encompass the effect of other environmental parameters, independent of bedstress, but still characteristic of the difference between the different types of habitats encompassed by the sampling stations: “Mediterranean offshore circalittoral mixed sediment” for stations E to J, “Mediterranean offshore circalittoral mud” for stations K to P, and “Mediterranean circalittoral coarse sediment” for stations A and C. For instance, the Lacaze-Duthiers canyon can represent a barrier to species connectivity between the two groups of stations, with station A and C at the west of the canyon, while all other stations are at the east. The difference between the stations from the two distinct benthic habitats was also observed in other analyses carried on at the same sites, based on the morphological identification of foraminifera, nematodes and macrofauna (Barras et al., 2025), confirming the clear trend observed here with eDNA.

It has been shown that communities from stable sediments are usually more affected by trawling than communities that have already been shaped by perturbations in their environment (Hiddink et al., 2006; Kaiser, 1998; Van Denderen et al., 2015; Watling and Norse, 1998). The marked hydrodynamic difference between the stations might therefore limit our power to detect the effect of trawling on the communities. A larger sampling in a habitat-homogenous area could therefore help to have a more precise description of the effect of the different temporal integrations of the trawling pressure on this large spectrum of benthic organisms (prokaryotes, protists, Animalia). Such sampling could allow detecting more subtle changes in the communities along the trawling gradient, such as a higher presence of scavengers in more trawled areas.

The global examination of the link between alpha diversity in the three taxonomic groups and the trawling pressure still enabled highlighting the pervasive effect of trawling on benthic organisms, even on the small and short-lived prokaryotes. Surprisingly, however, the station that was more intensively trawled in most time periods (the K station) did not display stark differences in terms of the alpha or beta diversity of the communities compared to other stations of its habitat group (stations O and P). Overall, trawling thus seemed to explain a modest part of the structure of the communities, possibly due to a threshold effect. The first occurrence of trawling is the one that may in fact have the largest impact on ecosystems: it was found to be more important in structuring the communities than the intensity of subsequent trawling events (Bonthond et al., 2023; Hiddink et al., 2006; Tinlin-Mackenzie et al., 2023). Additionally, a chronic trawling pressure will select for communities that are better adapted to this pressure (Hiddink et al., 2017), a phenomenon already demonstrated in our study area (Jac et al., 2022, 2020). The historical pervasive trawling activity on the whole continental shelf of the Gulf of Lion therefore complicates the impact assessment for this fishing technique, not only by the absence of untrawled control sites, but also by the ancient preselection of trawling-adapted organisms that are expected to display a marginal level of variation along the trawling gradient.

### 4.4. Conclusions

This cross-domain eDNA metabarcoding approach provided insights into the effect of natural and anthropic disturbances of the sediment on prokaryote, protist and Animalia communities. It demonstrates the relevance of eDNA approaches to comprehensively characterise benthic communities for impact studies, since it enables the evaluation of communities otherwise hardly accessible, such as prokaryotes and protists. Owing to their short generation time, small size and high metabolic plasticity, microorganisms can respond more rapidly to environmental disturbances than larger fauna, making them particularly informative indicators of early ecological changes. Furthermore, changes in microbial communities following bottom trawling may trigger cascading effects on the benthic ecosystem, for instance, by altering sediment biogeochemical processes and properties, with potential consequences for other organisms. Including microorganisms in impact assessments therefore provides a more comprehensive understanding of bottom trawling effects. We showed here that trawling pressure, integrated over different temporal scales, has an effect on the diversity of two of the three taxonomic groups considered. Interestingly, the main effect of trawling on microorganisms was actually a boosting effect on communities’ diversity, possibly due to resuspension induced by trawling, making the nutrients present in the sediment more accessible. This was also observed for Animalia, accompanied by a significant reduction of benthic fauna diversity with long-term trawling, showing the importance of the temporal scale considered when assessing the impact of trawling pressure. Our results also underline the need to sample and analyse data not only on the surface, but also in the deeper layers of sediment as stratification can be strongly affected by bottom trawling, and its analysis is thus an objective *per se*. In fact, the absence of vertical structure within the upper 15 cm, unusual for marine sediments, raises the hypothesis that repeated trawling homogenises the sediment column; testing this hypothesis would require layered sampling in comparable untrawled areas. The absence of such genuine trawl free area in the Gulf of Lion shelf, including in the Marine Park, is a cause for concern, and limits the proper assessment of the full extent of trawling impacts since the first event of bottom trawling. This calls for the establishment of trawl-free references that should serve as baseline areas.

## Credit authorship contribution statement

SAH, SV, C.Barras and C.L designed the sample collection, S.V collected the samples and prepared the abrasion and effort data with the help of Ifremer Système d’Information Halieutique, G.M provided sediment particle size data, S.A.H, C.Beaulieu and N.H ran the bioinformatic analyses, T.H conducted exploratory analyses, C.Beaulieu analysed the data and prepared all figures, C.Beaulieu, S.A.H and S.V wrote the original draft, C.Barras, C.L, M.S, J.R.O, A.P, G.M and N.H substantially contributed to revised drafts.

## Data availability

Sampling metadata and scripts are available at https://zenodo.org/records/22893857?preview=1&token=eyJhbGciOiJIUzUxMiJ9.eyJpZCI6ImI1OGM4MjdhLTNkOGEtNGExZi05NDg0LTk2NmIzY2IxZmRhOSIsImRhdGEiOnt9LCJyYW5kb20iOiIxNjY4YjVkMWZiYzRlMDE0ZmYzMDhiMWYwZTY0NTJhMyJ9.n_JV_tD0kCjsUtATIpIFufEMFyadCwpcARTUjtiaOqvMXXCfssL2kXHakX-4DRdr7uFdH5aIjCSwgJWYGdywYQ

## Acknowledgments

This project was funded by the OFB IMPEC project (OFB-20-844) and the PPR Ocean-Climate FUTURE-OBS project (ANR-22-POCE-0004), based on samples collected during the IMPEC cruise (LABRUNE Céline (2021) IMPEC cruise, RV L’Europe, https://doi.org/10.17600/18001594) onboard of the N/O l’Europe. We thank Ines Brian and Nathalie Ulysse for their contribution to DNA extractions, and the team of Argaly (https://www.argaly.com) for the production of metabarcode libraries.

## Supplementary data legends

**Supplementary Figure 1**, Information on the taxonomic assignment of the 16S marker, before the filtering of ASV, that kept only ASV assigned at least to the phylum level, to Bacteria or Archaea kingdoms. (A) Barplots of numbers of ASV and number of reads assigned up to a given taxonomic level for the 16S marker. There a lower proportion of unassigned in reads than in ASV, hinting at the fact that most assigned ASV are supported by a non-negligible number of reads. (B) Taxonomic assignment of ASV in major prokaryote phylum

**Supplementary Figure 2**, Information on the taxonomic assignment of the 18S-V9 marker, before (A, B) and after (C) the filtering of ASV to retain ASV assigned to a protist class. (A) Barplots of numbers of ASV and number of reads assigned up to a given taxonomic level for the 18S-V9 marker. There a lower proportion of unassigned in reads than in ASV, hinting at the fact that most assigned ASV are supported by a non-negligible number of reads. (B) Proportions of ASV assigned to protists, other organisms or non-assigned at the class level. (C) Taxonomic assignment of ASV in major protist Subdivisions, after filtration to keep only ASV belonging to protist class

**Supplementary Figure 3**, (A) Categories of taxonomic assignment of OTU for the 4 markers used to detect Animalia, before any filtering for Animalia. The categories are based on the phylum assignment of the OTUs. The COI marker is the one that has the largest proportion of Animalia OTUs but also the lower number of OTU overall. 18S-V1 and V4 have a lot of OTUs assigned to organisms that are not Animalia (protists for instance). 18S-V9 has a lot of unassigned OTUs and detects a small proportion of Animalia among the numerous OTUs it detects. (B) Barplots of numbers of OTU assigned up to a given taxonomic level for the 4 markers used to detect Animalia, before the filtering. The COI marker is the most efficient to assign up to precise taxonomic ranks (most OTUs are assigned up to the species), and 18S markers also mostly assign up to the class or order. (C) Major Animalia phyla detected by each marker after the filtering steps (keep only Animalia, discard terrestrial Insecta, Arachnida, Mammalia and Aves). Some phyla are only identified thanks to the COI marker, such as the Mollusca and Porifera, but Nematoda, Platyhelminthes and Xenacoelomorpha are better detected with the 18S markers, which shows well the complementarity of using multiple markers to get the full scope of benthic animalia. Nematoda are the most abundant phyla detected. (D) Barplots of numbers of OTU assigned up to a given taxonomic level for the 4 markers used to detect Animalia, after the filtering. Almost all COI OTU are assigned up to the species, and most OTU from 18S markers are assigned up to the order.

**Supplementary Figure 4**, Rarefaction curves showing the evolution of the number of ASV detected depending on the number of reads, for each sample. The curve for each sample is coloured according to the sediment layer where the sample comes from, and grouped by station (A.I) Rarefaction curves with the 16S marker for prokaryotes. (A.II) Zoom on the rarefaction curves of prokaryotes for samples with low number of reads (under 100 000 reads). (B.I) Rarefaction curves with the 18S-V9 marker for protists. (B.II) Zoom on the rarefaction curves of protists for samples with low number of reads (under 100 000 reads).

**Supplementary Figure 5**, Results of the ordinations on the communities of Animalia (A), protists (B) and prokaryotes (C). (I) shows the PCoA of the different stations and horizons, from a Bray Curtis distance matrix for prokaryote and protist, or a Jaccard distance matrix for Animalia. Ellipses are drawn to encompass the points from the same station. Environmental variables are fitted onto the ordination with the envfit function from vegan, and the darker arrows are the variables kept in a forward selection of variables in a distance-based RDA. (II) Same ordination, with ellipses drawn around same horizons of the sediment. No trend can be seen from the structure of the communities in the different layers of sediment. (III) Partition of the variation in community distance matrices, based on trawling (SAR_P90, SAR_wmean_2019_2021, SAR_072021, EP_h_per_month, EP_h_per_week) and natural (bedstress) explanatory variables. Very little of the distance between communities in the samples is explained by these variables.

**Supplementary Figure 6**, Overview of the sediments layers community content in benthic and pelagic organisms in all stations, for Animalia (A) and protists (B). The benthic or pelagic lifestyle of each protist ASV or Animalia OTU was extracted from the WoRMS database, based on their taxonomic assignment. For protists a lot of ASV could not be assigned to the benthic or pelagic lifestyle, while in Animalia most OTU detected in the sediments appear to belong to the benthic compartment.

**Supplementary Figure 7**, Overview of the trends between the alpha diversity of organisms and the trawling and bedstress variables, the blue line being the outcome of a lm between each pair of variables. For Animalia (A), only benthic OTUs were taken for the calculation of the alpha diversity, while for protists (B) and prokaryotes (C), all ASV were used to calculate the Shannon diversity.

**Supplementary Table 1**, Numeric values of environmental variables or trawling pressure variables for all stations and sediment layers.

**Supplementary Table 2**, Details about the different software versions and sets of parameters used in the bioinformatic treatment pipeline that was used to process the metabarcoding data.

**Supplementary Table 3**, List of species typical from the Gulf of Lion identified in literature, with the links for the associated papers mentioning them.

**Supplementary Table 4** (Sheet 1) Tracking of the number of reads for different steps of the bioinformatic treatment pipeline, for the 5 different markers. (Sheet 2) Tracking of the number of OTU along the different filtration steps, in the 4 markers used for Animalia. (Sheet 3) Tracking of the number of ASV along the different filtration steps, in the 18S-V9 marker used for protists. (Sheet 4) Tracking of the number of ASV along the different filtration steps, in the 16S marker used for prokaryote.

**Supplementary Table 5**, Additional information on some generalised linear mixed models.

(Sheet 1) Details of the results of the modelling of the link between the alpha diversity of Animalia (number of OTU) against the alpha diversity of protists and prokaryotes (Shannon index and Pielou’s J for eveness), but also of the alpha diversity of benthic Animalia only, against the same indexes, and of the alpha diversity of protists against the one of prokaryotes. The estimates and p-values represented here are from glm on a negative binomial distribution for Animalia, and for a lm on the transformed Shannon index for protist. R^2^ of the models are given, highlighting how much of the variability of the alpha diversity is explained by the predictors. We can see that alpha diversity in protists can explain most of the alpha diversity in Animalia, but that alpha diversity in prokaryotes doesn’t explain much of the alpha diversity in protists. (Sheet 2) Hierarchical partitioning on the model linking environmental predictors to the granulometry. We can see that that bedstress accounts for 35% of the 88% of the variance of granulometry explained by the model. Other variables that explain a non-negligible amount of the variance of granulometry are water depth (22.8%), and swept area ratio variables: during the past month (SAR_07_2021: 9.5% of the variance explained by the model), during the past 3 years (SAR_wmean_2019_2021: 12.42% of the variance) and during the past 10 years (SAR_P90: 11.6% of the variance).

