## Supplementary figures and images for "Pervasive effect of bottom trawling on benthic biodiversity unravelled by metabarcoding across the Tree of Life"

### Supplementary Figure 1

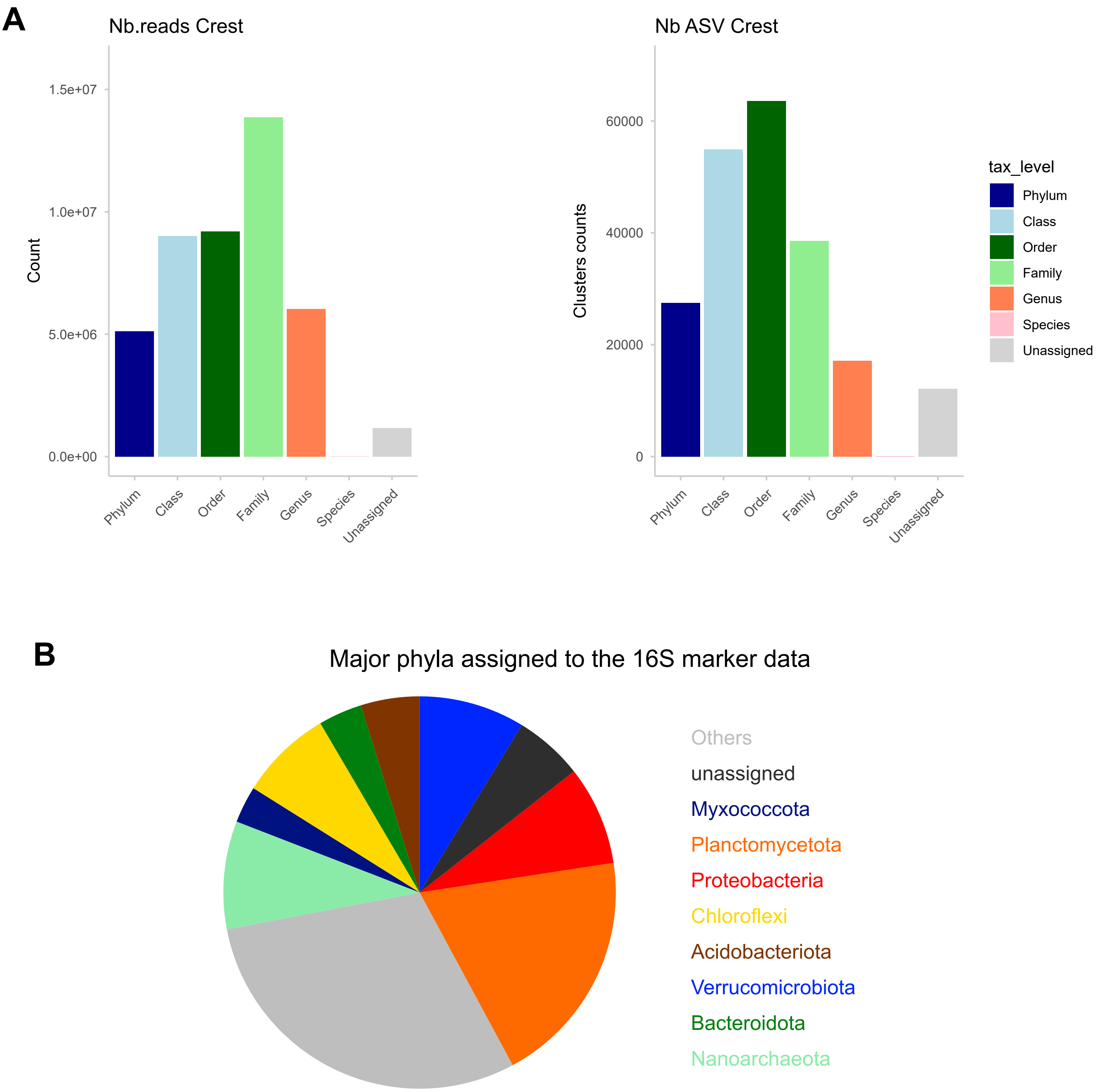

### Supplementary Figure 3

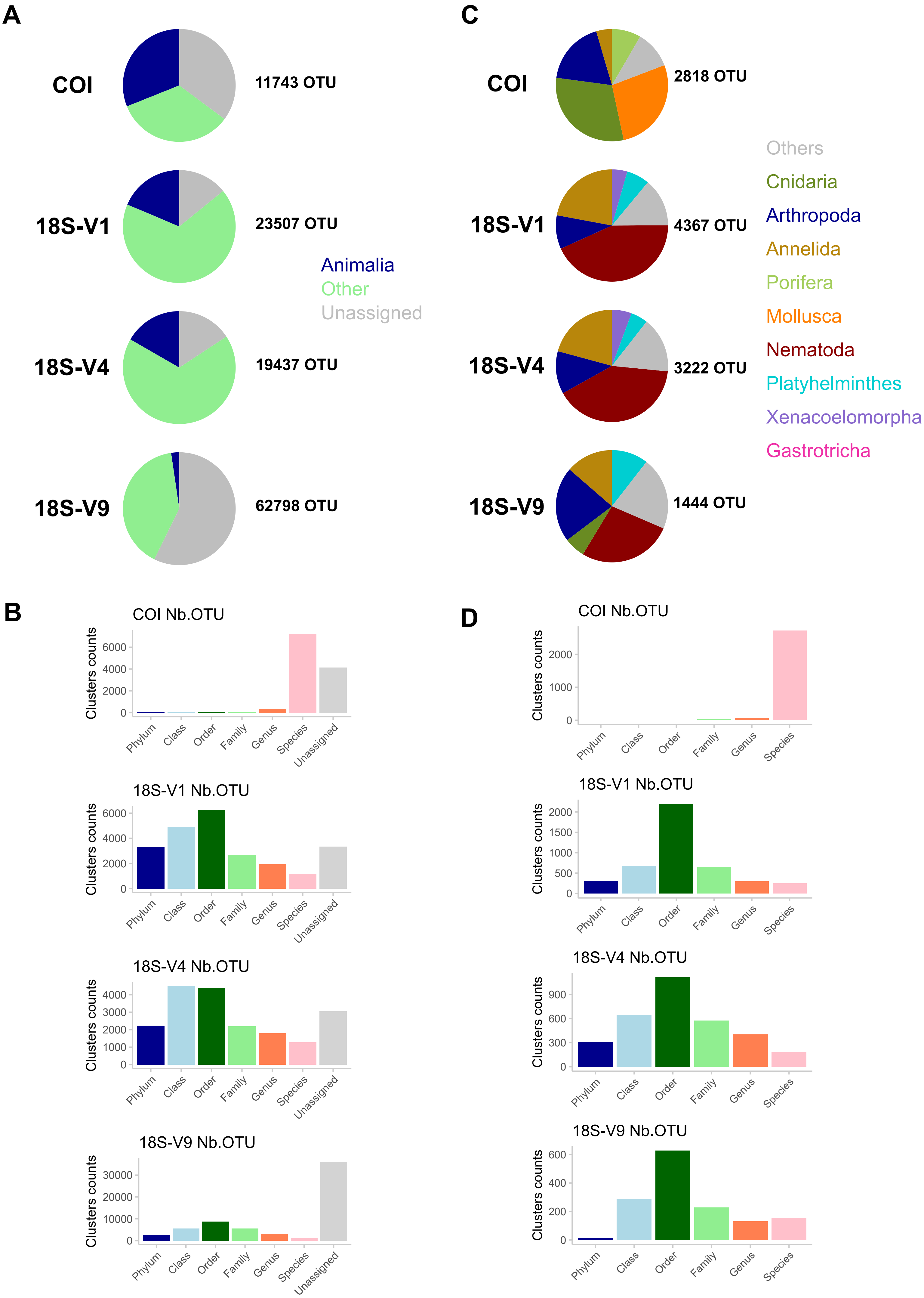

### Supplementary Figure 4

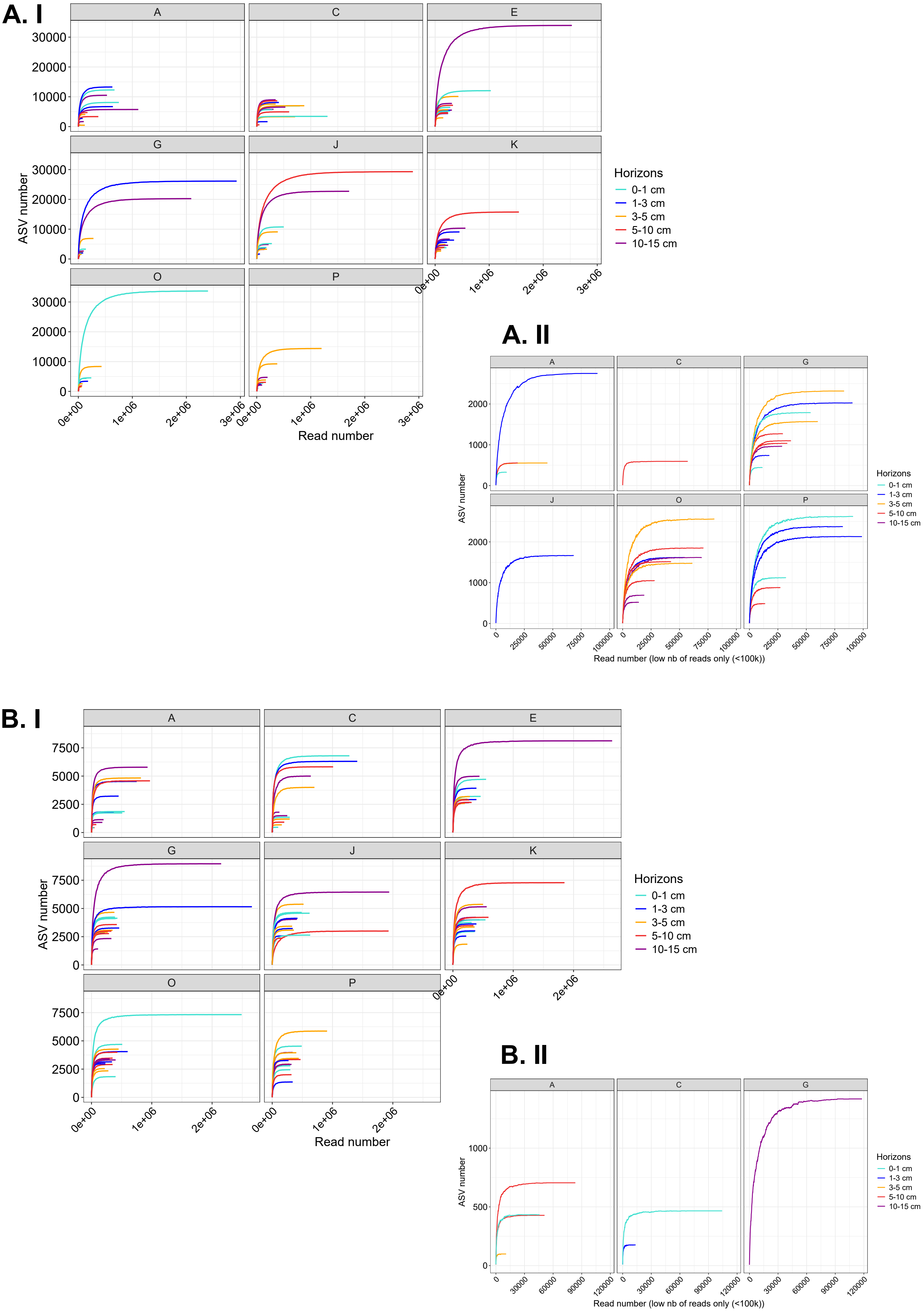

### Supplementary Figure 5

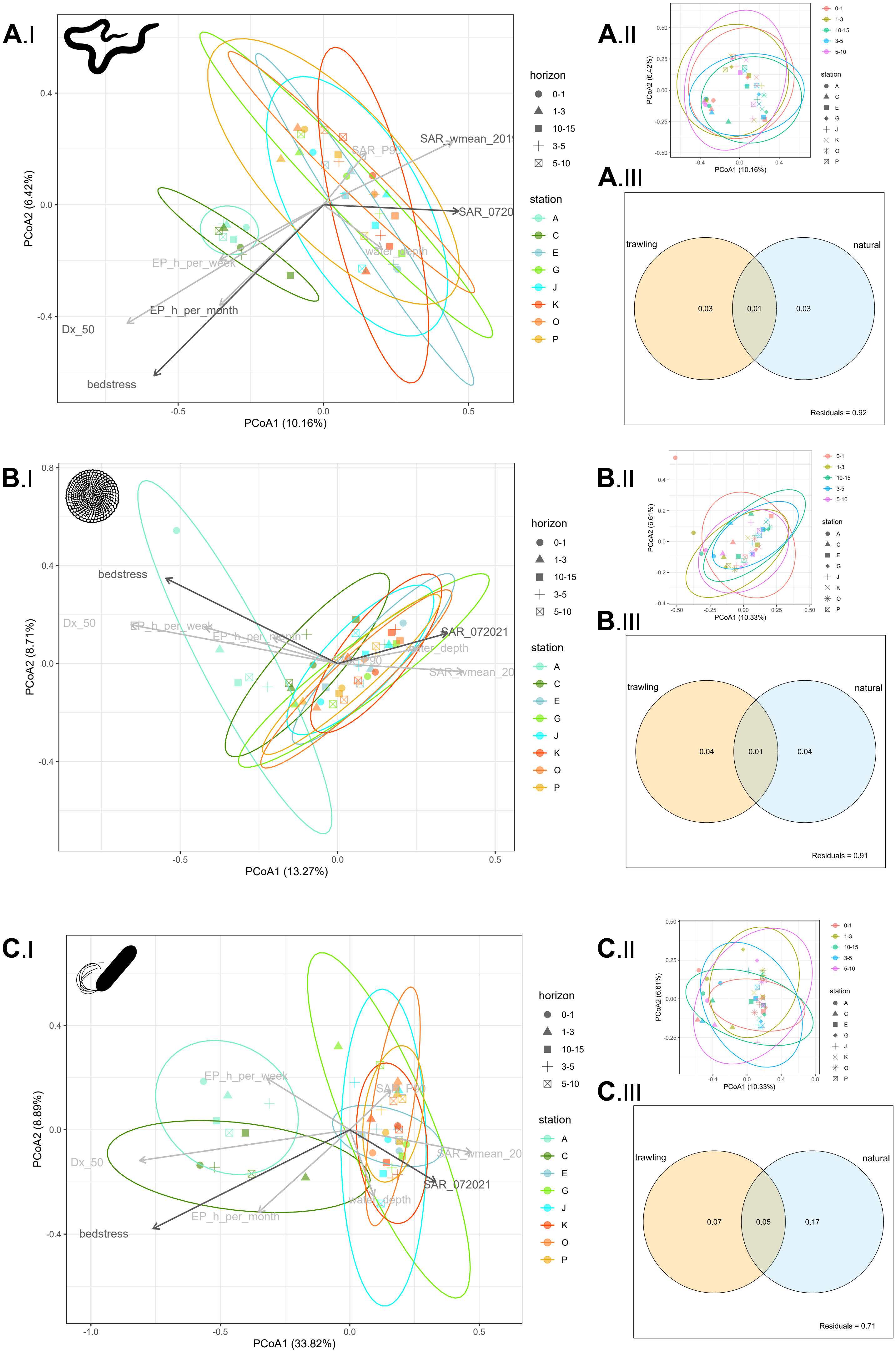

### Supplementary Figure 6

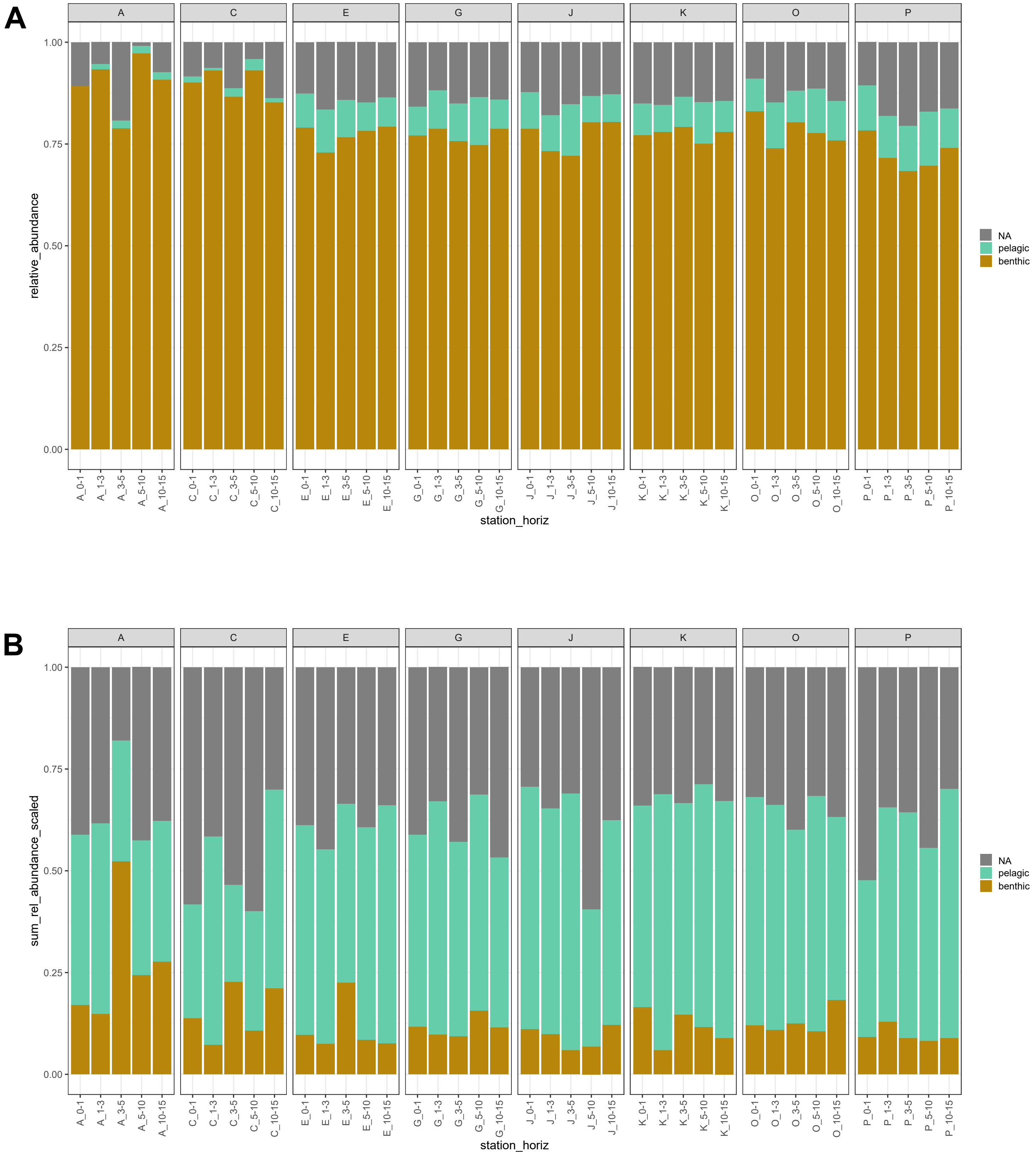

### Supplementary Figure 7

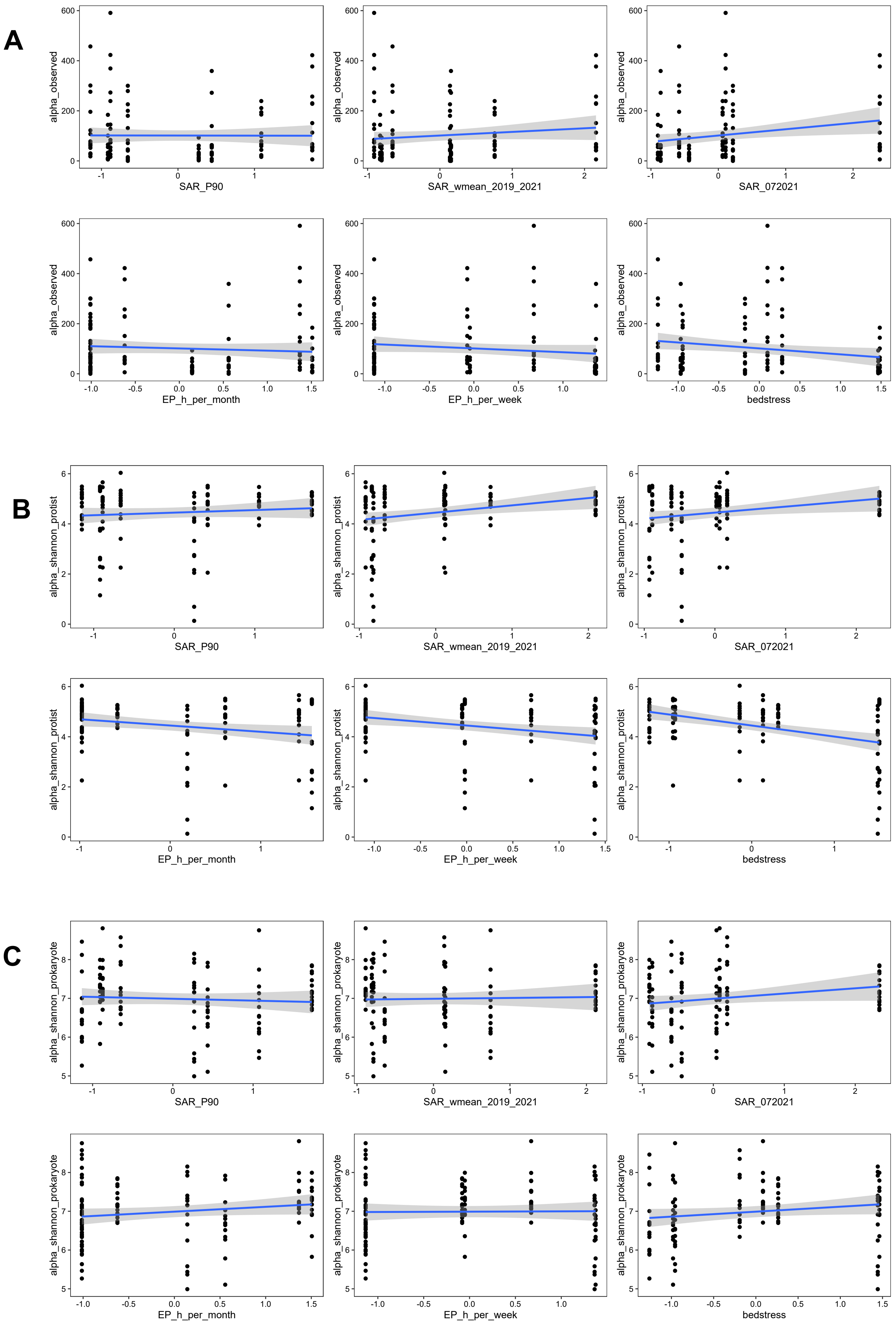
