## Supplementary Figure 2 for "Pervasive effect of bottom trawling on benthic biodiversity unravelled by metabarcoding across the Tree of Life"

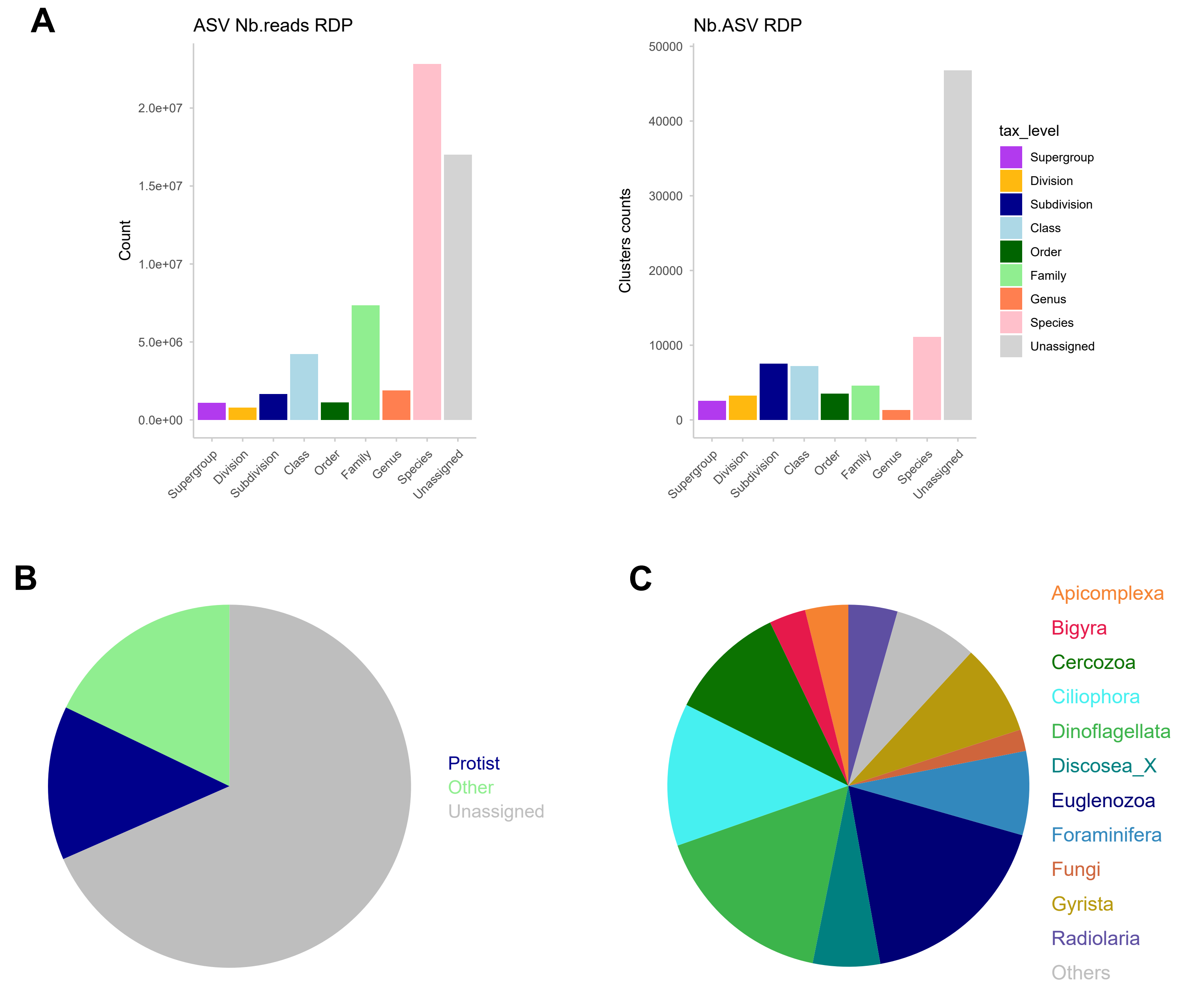


Categories of ASV taxonomic assignment, for 18S-V9 marker, based on class (87 770 ASV in total)

Major Subdivisions assigned to the 18S-V9 marker, in protist-filtered dataset (10 306 ASV)
